# MaBoSS for HPC environments: Implementations of the continuous time Boolean model simulator for large CPU clusters and GPU accelerators

**DOI:** 10.1101/2024.03.18.585487

**Authors:** Adam Šmelko, Miroslav Kratochvíl, Emmanuel Barillot, Vincent Noël

## Abstract

Computational models in systems biology are becoming more important with the advancement of experimental techniques to query the mechanistic details responsible for leading to phenotypes of interest. In particular, Boolean models are well fit to describe the complexity of signaling networks while being simple enough to scale to a very large number of components. With the advance of Boolean model inference techniques, the field is transforming from an artisanal way of building models of moderate size to a more automatized one, leading to very large models. In this context, adapting the simulation software for such increases in complexity is crucial. We present two new developments in the continuous time Boolean simulators: Ma-BoSS.MPI, a parallel implementation of MaBoSS which can exploit the computational power of very large CPU clusters, and MaBoSS.GPU, which can use GPU accelerators to perform these simulations.

## Introduction

Biological systems are large and complex, and understanding their internal behavior remains critical for designing new therapies for complex diseases such as cancer. A crucial approach in this endeavor is building computational models from existing knowledge and analyzing them to find intervention points and to predict the efficacy of new treatments. Many different frameworks have been used to describe biological systems, from quantitative systems of differential equations to more qualitative approaches such as Boolean models. While the former seems more adapted to represent complex behavior, such as non-linear dependencies, the latter is being increasingly used because of its capability to analyze very large systems. Many Boolean models have been built to describe biological systems to tackle a variety of problems: from understanding fundamental properties of cell cycle (1, 2) to advanced properties of cancer (3, 4).

Historically, the task of building Boolean models involved reading an extensive amount of literature and summarizing it in a list of essential components and their interactions. More recently, thanks to advances in databases listing such interactions (5, 6) and to experimental techniques providing information on a bigger number of components, the automatic methods have been designed to infer Boolean formulas from the constraints encoded in the knowledge and the experimental data (7–9), allowing construction of large Boolean models. While this effort faces many challenges, we believe it is a promising way to study the large-scale complexity of biological systems. However, in order to analyze the dynamic properties of such large Boolean models, we need to develop efficiently scalable simulation tools.

Here, we present adaptations of MaBoSS (10, 11) — a stochastic Boolean simulator that performs estimations of state probability trajectories based on Markov chains – to modern HPC computing architectures, which provide significant speedups of the computation, thus allowing scrutinization and analysis of much larger boolean models. The main contributions comprise two new implementations of MaBoSS:

- MaBoSS.GPU, a GPU-accelerated implementation of MaBoSS, which is designed to exploit the computational power of massively parallel GPU hardware.
- MaBoSS.MPI, a parallel implementation of MaBoSS which can scale to multinode environments, such as large CPU clusters.

The source code of the proposed implementations is publicly available at their respective GitHub repositories^1 2^. We also provide the scripts, presented plots, data and instructions to reproduce the benchmarks in the replication package^3^.

To showcase the utility of the new implementations, we performed benchmarking on both existing models and large-scale synthetic models. As the main results, MaBoSS.GPU provided over 200× speedup over the current version of Ma-BoSS on a wide range of models using contemporary GPU accelerators, and MaBoSS.MPI is capable of almost linear performance scaling with added HPC resources, allowing similar speedups by utilizing the current HPC infrastructures.

## Background

### Boolean signaling models

A Boolean signaling model consists of *n* nodes which are either active or inactive, gaining values 1 or 0 respectively. The *state* of the whole model is represented by a vector *S* of *n* Boolean values where *S*_*i*_ represents the value of the *i*-th node. We denote the set of all possible states as *𝒮* = {0, 1}^*n*^; thus |*𝒮*| = 2^*n*^.

Interactions in the model are described as transitions between two states. A single state can have multiple transitions to other states with assigned transition probabilities. In turn, a Boolean network is represented as a directed weighted graph *G* = (*𝒮, ρ*), where *ρ* : *𝒮 ×𝒮 →* [0, *∞*) is a transition function generating *transition rates*. For convenience, it holds that:

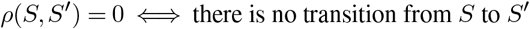

MaBoSS simulates the *asynchronous update strategy*, where at most a single node changes its value in each transition — consequently, a state *S* can have at most *n* possible transitions.

To determine the possible transitions, each node follows the *Boolean logic ℬ*_*i*_ : *𝒮 →* [0, *∞*), which determines the expected Poisson-process rate of transitioning to the other value. If *ℬ*_*i*_(*S*) = 0, then the transition at node *i* is not allowed in state *S*. Given this formalization, the simulation can be also viewed as a continuous-time Markov process.

MaBoSS algorithm simulates the above process to produce stochastic *trajectories*: sequences of states *S*^0^, *S*^1^, …, *S*^*k*^ and time points *t*^0^ *< t*^1^ *< ···< t*^*k*^ where *t*^0^ = 0 and *S*^0^ is the initial state, and for each *i* ∈ {0,…, *k −* 1}, *S*^*i*^ transitions to *S*^*i*+1^ at time *t*^*i*+1^. The simulation ends either by a timeout when reaching the maximal allowed time, or by reaching a fixed point state with no outgoing transitions. The algorithm for a single iteration of the trajectory simulation is given explicitly in Algorithm 1.

#### Algorithm 1

A single iteration of the MaBoSS simulation of a trajectory, given the state *S* and time *t*.

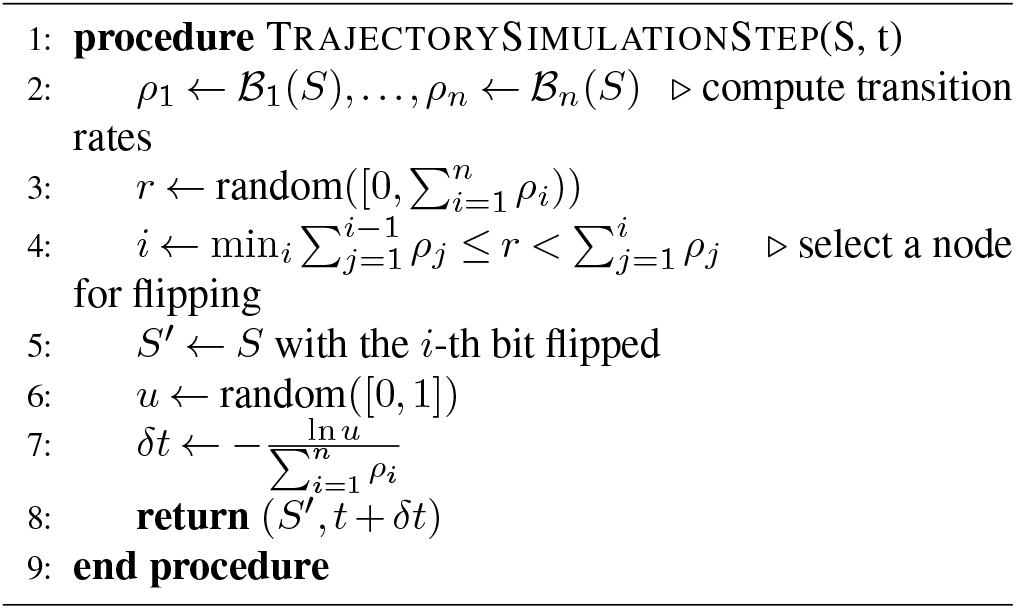

To obtain a good overview of the Markovian process, multiple trajectories are generated and aggregated in compound trajectory statistics. Commonly obtained statistics include:

#### Network state probabilities on a time window

Trajectory states are divided by their transition times into time windows based on the time intervals specified by a window size. For each window, the probability of each state is computed as the duration spent in the state divided by the window size. The probabilities of the corresponding windows are then averaged across all sub-trajectories.

#### Final states

The last sampled states from the trajectories are used to compute a final state distribution.

#### Fixed states

All reached fixed points are used to compute a fixed state distribution.

To maintain the brevity in the statistics, MaBoSS additionally allows marking some nodes *internal*. This is useful because nodes that are not “interesting” from the point of final result view occur quite frequently in Boolean models, and removing them from statistics computation often saves a significant amount of resources.

### Computational complexity of parallel MaBoSS algorithm

#### Simulation complexity

We estimate the time required to simulate *c* trajectories as follows: For simplification, we assume that a typical Boolean logic formula in a model of *n* nodes can be evaluated in *𝒪* (*n*) (this is a very optimistic but empirically valid estimate). With that, the computation of all possible transition rates (Algorithm 1, line 2) can be finished in *𝒪* (*n*^2^). The selection of the flipping bit (Algorithm 1, line 4) can be finished in *𝒪* (*n*), and all other parts of the iteration can finish in *𝒪* (1). In total, the time complexity of one iteration is *𝒪* (*n*^2^). If we simulate *c* trajectories with an upper bound of trajectory length *u*, the simulation time is in *𝒪* (*c ·u ·n*^2^).

In an idealized PRAM model with infinite parallelism, we can optimize the algorithm in the following ways:

- Given *c* processors, all trajectory simulations can be performed in parallel, reducing the time complexity to *𝒪* (*m ·n*^2^). (Note that this does not include the results aggregation. See *Statistics aggregation* section for further description.)
- With *n* processors, the computation of transition rates in the simulation can be done *𝒪* (*n*) time, and the selection of the flipping bit can be done in *𝒪* (log *n*) time using a parallel prefix sum, giving *𝒪* (*n*) time for a single iteration.

Thus, using a perfect parallel machine with *c ·n* processors, the computation time can be reduced to *𝒪* (*u ·n*). Notably, the *𝒪* (*u*) simulation steps that must be performed serially remain a major factor in the whole computation time.

#### Statistics aggregation

The aggregation of the statistics from the simulations is typically done by updating a shared associative structure indexed by model states, differing only in update frequency between the three kinds of collected statistics.

If the associative structure is implemented as a hashmap, the updates can be done in *𝒪* (1) for a single process. With multiple processors, the algorithm may hold partial versions of the hashmap for each processor, and aggregate all of them at the end of the computation, which can be done in *𝒪* (log *c ·m*) using *c* processors, assuming the maximal size of statistic to be *m*.

As an interesting detail, the hash structures pose a surprising constant-factor overhead. In networks where most nodes are internal, the hash map may be replaced by a fixed-size multidimensional array that holds an element for all possible combinations of external node values (basically forming a multidimensional histogram). We discuss the impact of this optimization in the next section.

## Implementation

### MaBoSS.GPU

#### Simulation

In the CPU version of MaBoSS, the simulation part is the most computationally demanding part, with up to 80% of MaBoSS runtime spent by just evaluating the Boolean formulas (the exact number depends on the model). The original formula evaluation algorithm in MaBoSS used a recursive traversal of the expression tree, which (apart from other issues) causes memory usage patterns unsuitable for GPUs: the memory required per each core is not achievable in current GPUs, and there are typically too many cache misses (12).

There are multiple ways to optimize the expression trees for GPUs: One may use a linked data structure that is more cache-friendly such as the van Emde Boas tree layout (13), or perhaps represent the Boolean formulas as a compact continuous array, or convert it to CNF or DNF bitmasks that can be easily evaluated by vector instructions. We decided to leave the exact representation choice on the compiler, by encoding the expressions as direct code and using the runtime compilation of GPU code (14). Using this technique, the Boolean formulas are compiled as functions into a native binary code, which is directly executed by the GPU. As the main advantage, the formulae are encoded in the instructions, preventing unnecessary fetches of the encoded formulas from other memory. At the same time, the compiler may apply a vast spectrum of optimizations on the Boolean formulas, including case analysis and shortcutting, again resulting in faster evaluation.

A possible drawback of the runtime compilation stems from the relative slowness of the compiler — for small models, the total execution time of MaBoSS.GPU may be easily dominated by the compilation.

Also, due to the involved implementation complexity, we avoided optimization of the computation of individual trajectories by splitting the Boolean function evaluation into multiple threads (thus missing the factor of *n* threads from the asymptotic analysis). While such optimization might alleviate some cache pressure and thus provide significant performance improvements, we leave its exploration to future work.

#### Statistics aggregation

For optimizing the statistics aggregation, MaBoSS.GPU heavily relies on the fact that the typical number of non-internal nodes in a real-world MaBoSS model rarely exceeds 10 nodes, regardless of the size of the model. This relatively low number of states generated by non-internal nodes allows us to materialize the whole statistics structure (called “histogram”) as a fixed-size array (rarely exceeding 2^10^ elements).

This approach allows us to avoid storing the states as the keys, and gives a simple approach that can map the state to the histogram index using simple bit masking and shifting instructions. Further, we use several well-known GPU histogram update optimizations to improve the performance, including shared memory privatization and atomic operations.

### MaBoSS.MPI

MaBoSS.MPI is a straightforward extension of the original MaBoSS CPU code to the MPI programming interface. Briefly, each thread is assigned to compute a single trajectory at once, progressively collecting the results into a privatized hashmap-based statistics aggregation structure. The total amount of trajectory tasks is evenly split among the MPI nodes, adding a second layer of parallelism.

Once all trajectory simulations are finished and the statistics are computed for each thread, the intermediate data are reduced into the final result using MPI collective operations.

## Results

To evaluate the impact of the implemented optimizations, we present the results of performance benchmarks for Ma-BoSS.GPU and MaBoSS.MPI by comparing their runtimes against the original CPU implementation. To obtain a comprehensive overview of achievable results, we used both real-world models and synthetic models with varying sizes.

### Benchmarking Methodology

For the benchmarks, we used 3 real-world models of 10, 87 and 133 nodes (cellcycle (1), sizek (2) and Montagud (4)). In order to test the scalability of the GPU and MPI implementation, we also created several synthetic models with up to 1000 nodes. Synthetic models were designed in a way such that the length of each trajectory is predictable, and the models have no stable states. Also, the number of non-internal nodes was kept low (10 nodes) to enable the usage of the histogram optimization. The synthetic models together with their Python generator are available in the replication package.

The GPU implementation benchmarks were run on a datacenter-grade NVIDIA Tesla A100 GPU and a consumer-grade NVIDIA RTX 3070 Laptop GPU. The CPU implementation benchmarks were run on a 32-core Intel Xeon Gold 6130 CPU with multithreading. The CPU implementation was compiled with GCC 13.2.0, and the GPU implementation was compiled with CUDA 12.2. Each measurement was repeated 10 times, and the average runtime was used as the final result.

The MPI implementation benchmarks were run on the MareNostrum 4 supercomputer ^4^.

### Performance of MaBoSS.GPU

In Figure 1, we compare the wall time of the CPU and GPU implementations on real-world datasets. The GPU implementation is faster than the CPU implementation on all models, and the speedup shows to be more significant on the models with more nodes and longer trajectories. On the MONTAGUD model with 133 nodes, but a relatively short average trajectory, we achieve 145*×* speedup. On a slightly smaller SIZEK model with a longer average trajectory, the speedup is up to 326*×*.

**Fig. 1.**
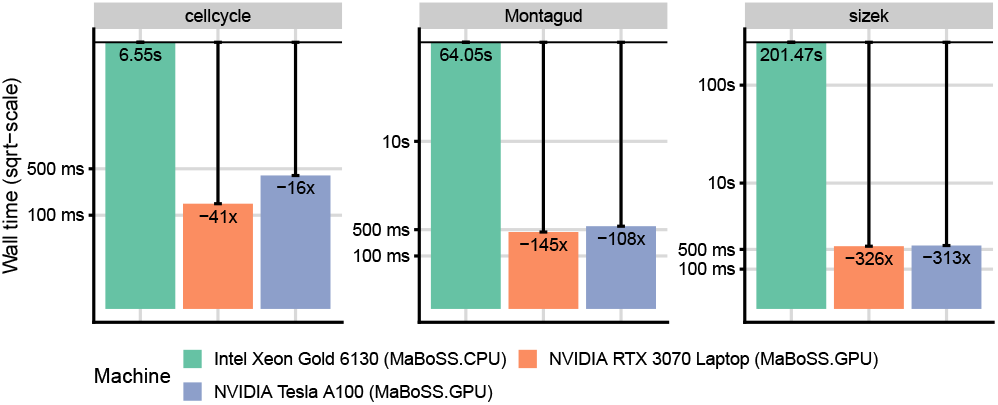
Wall time comparison of MaBoSS and MaBoSS.GPU on real-world models. Each model is simulated with 1 million trajectories.

Figure 2 shows much finer performance progression on synthetic models. We observed that the CPU variant starts to progress steeper at around the 100 nodes boundary. We assume that the implementation hits the cache size limit, and the overhead of fetching the required data from the memory becomes dominant. The same can be observed in the GPU variant later at around 200 nodes. Expectably, the cache-spilling performance penalty is much more significant on GPUs. Overall, the results suggest that the optimization of dividing transition rate computations among multiple threads, as mentioned in *Implementation* section, may provide a better speedup for bigger models, as it alleviates the register and cache pressure.

**Fig. 2.**
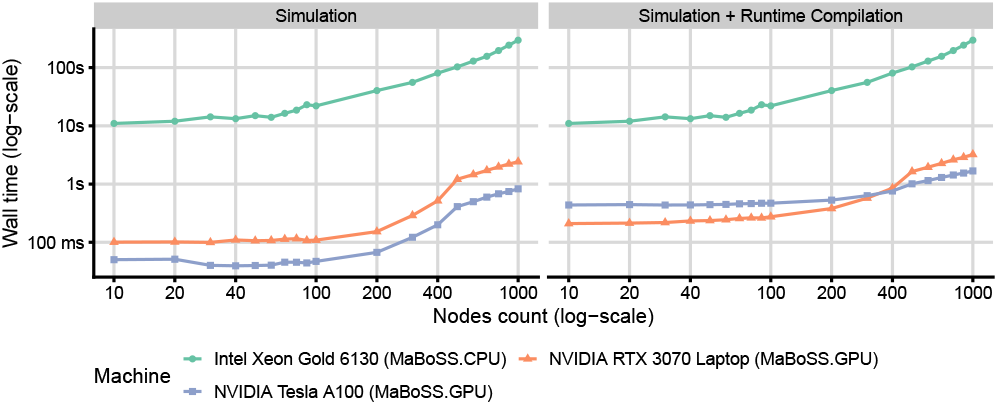
Wall time comparison of MaBoSS and MaBoSS.GPU on synthetic models. Each model is simulated with 1 million trajectories. The two panels differ by the inclusion of the runtime GPU code compilation, showing its impact on total run time.

Additionally, Figure 2 shows the total runtime of the GPU implementation including the runtime compilation step. Comparing the panels, we observe that the relative run-time compilation overhead quickly disappears with increasing model size. Figure 3 shows the results of more detailed benchmarks for this scenario, as run on the NVIDIA Tesla A100 GPU. We observed that the compilation time is linearly dependent on the number of nodes and formula lengths (measured in the number of occurring nodes). Notably, as soon as the simulation becomes more complex (e.g., by increasing the number of nodes or simulated trajectories), the compilation time becomes relatively negligible even for models with unrealistically long formulae. This suggests that the runtime compilation is a viable optimization methodology also for much larger models.

**Fig. 3.**
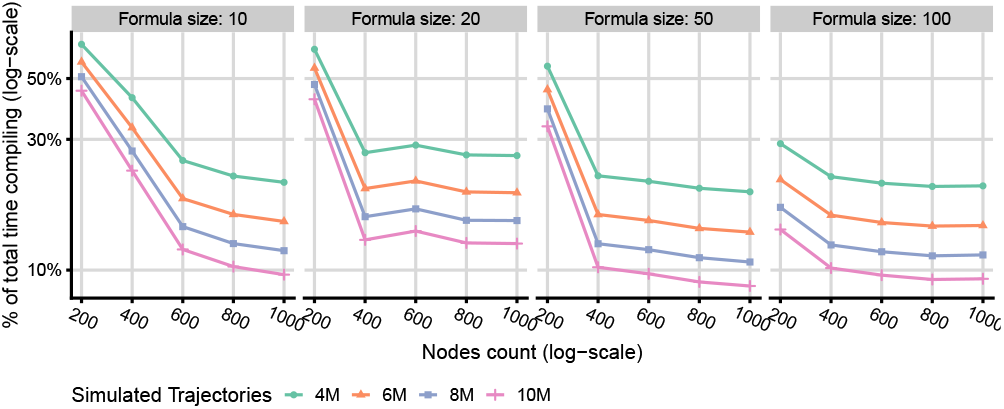
The time spent in the runtime compilation of the Boolean formulas simulating models with varying numbers of nodes, trajectories, and formula lengths.

### Performance of MaBoSS.MPI

Figure 4 shows the efficiency of the MaBoSS.MPI implementation on the SIZEK model. We ran multiple suites, ranging from a single MPI node up to 192 nodes, each running 20 cores. We can observe a close-to-linear speedup of up to 64 MPI nodes (1280 cores), and a plateau for larger suites (Figure 4, green). This can be explained by hitting an expectable bottleneck in parallelization overhead and MPI communication cost when the problem is divided into too many small parts.

**Fig. 4.**
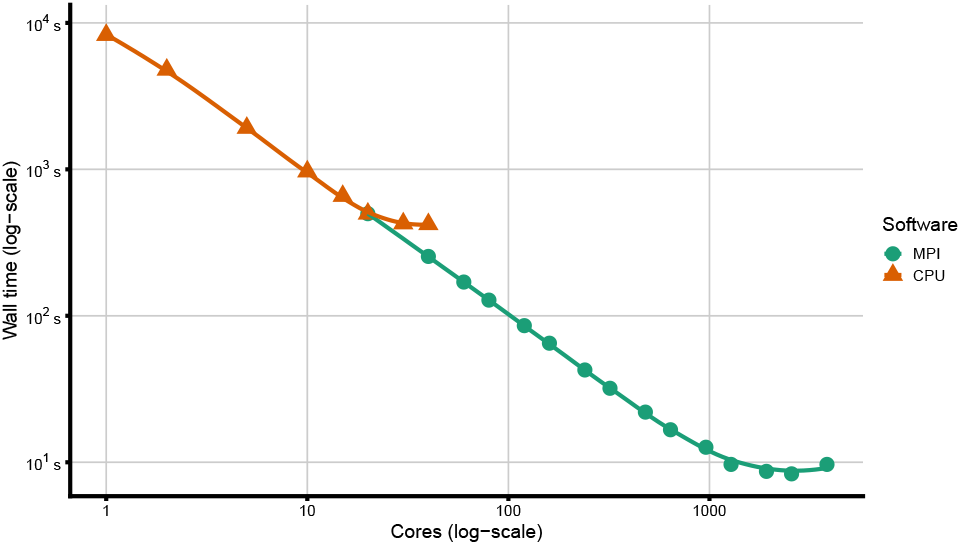
Scalability results of MPI implementation on Sizek model with 20 cores per MPI node.

To stress the scalability of the implementation, we also used a synthetic model with 1000 nodes. We simulated this model on 32 cores per MPI node, on 1 to 192 nodes (32 to 6144 cores). The obtained speedups are summarized in Figure 5.

**Fig. 5.**
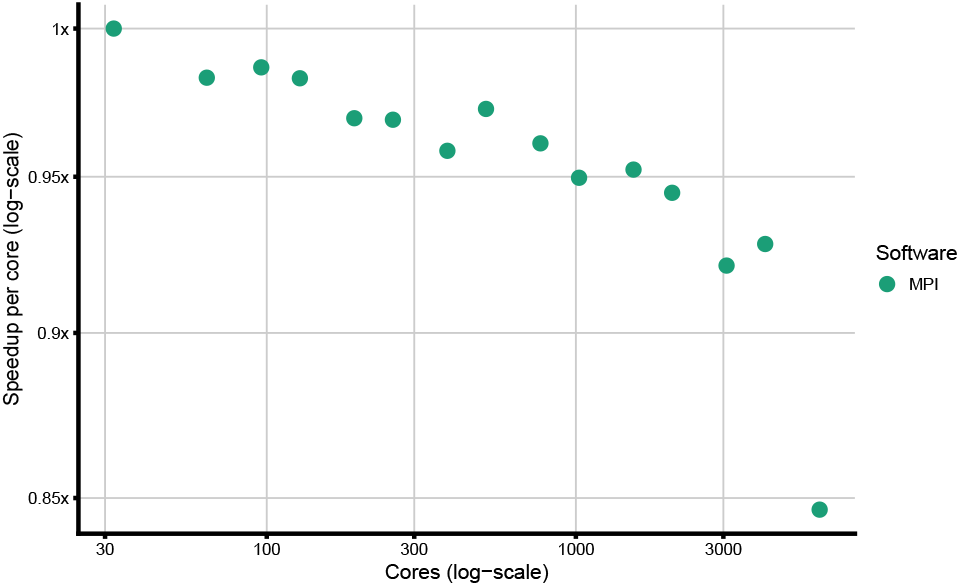
Speedup scaling of MPI implementation on a synthetic model with 1000 nodes and 32 cores per MPI node.

Using this configuration, the simulation time decreases from 20 hours on 1 MPI node to 430 seconds on 192 nodes. As expected, the plateau in the speedup was not observed in simulations that involve larger models.

## Conclusions

In this work, we presented two new implementations of Ma-BoSS tool, a continuous time Boolean model simulator, both of which are designed to enable utilization of the HPC computing resources: MaBoSS.GPU is designed to exploit the computational power of massively parallel GPU hardware, and MaBoSS.MPI enables MaBoSS to scale to many nodes of HPC clusters via the MPI framework. We evaluated the performance of these implementations on real-world and synthetic models and demonstrated that both variants are capable of providing significant speedups over the original CPU code. The GPU implementation shows 145–326× speedup on real-world models, and the MPI implementation delivers a close-to-linear strong scaling on big models.

Overall, we believe that the new MaBoSS implementations enable simulation and exploration of the behavior of very large, automatically generated models, thus becoming a valuable analysis tool for the systems biology community.

### Future work

During the development, we identified several optimization directions that could be taken by researchers to further scale up the MaBoSS simulation approach.

Mainly, the parallelization scheme used in MaBoSS.GPU could be enhanced to also parallelize over the evaluation of Boolean formulas. To avoid GPU thread divergence, this would however require a specialized Boolean formula representation, entirely different from the current version of Ma-BoSS; likely even denying the relative efficiency of the use of runtime compilation. On the other hand, this optimization might decrease the register pressure created by holding the state data, and thus increase the performance on models with thousands of nodes.

In the long term, easier optimization paths might lead to sufficiently good results: For example, backporting the GPU implementation improvements back to the MaBoSS CPU implementation could improve the performance even on systems where GPU accelerators are not available. Similarly, both MaBoSS.GPU and MaBoSS.MPI could be combined into a single software that executes distributed GPU-based analysis over multiple MPI nodes, giving a single high-performance solution for extremely large problems.

## ACKNOWLEDGEMENTS

We thank Laurence Calzone and Gautier Stoll for their guidance and fruitful discussions.

## Footnotes

1 https://github.com/sysbio-curie/MaBoSS.GPU

2 https://github.com/sysbio-curie/MaBoSS-env-2.0

3 https://github.com/asmelko/gigascience24-artifact

4 https://www.bsc.es/marenostrum/marenostrum

